## Supplementary material for "Differential nucleosome organization in human interphase and metaphase chromosomes": combined suppl files

#### Table of contents

2-3 Appendix Supplementary Figure Legend
4 Appendix Table S1
5 Apendix Table S2, Apeendix Table S3
6 Apendix Figure S1
7 Apendix Figure S2
8 Apendix Figure S3
9 Apendix Figure S4
10 Apendix Figure S5
11 Apendix Figure S6
12 Apendix Figure S7
13 Apendix Figure S8
14 Apendix Figure S9

**Appendix Supplementary Figure Legend**

**Appendix Figure S1. ECDF plots of local nucleosome occupancy correlation between interphase and metaphase across 18 chromatin states.** Empirical cumulative distribution function (ECDF) curves illustrate the local correlation in nucleosome occupancy between interphase and metaphase across 18 chromatin states defined by ChromHMM in HeLa S3 cells. ECDF curves for chromatin states E1, E2, E3, E4, E8, E9, E10, and E11 lie well above the genome-wide curve, indicating a marked enrichment of regions with low interphase–metaphase correlation.

**Appendix Figure S2. Linker DNA length distribution in interphase and metaphase across chromatin states.** Distribution of linker lengths inferred from unique nucleosome maps in interphase and metaphase across 18 chromatin states defined by ChromHMM in HeLa S3 cells.

**Appendix Figure S3. Genome-wide comparison of interphase and metaphase chemical nucleosome maps from clone-2 and clone 1-2.** (**A**) Empirical cumulative distribution function (ECDF) plots showing genome-wide local Pearson correlation of interphase and metaphase nucleosome occupancy scores between two H4S47C-expressing cell lines, clone-2 and clone 1-2. Correlations were calculated using a 501-bp sliding window with a 1-bp step size. (**B**) Frequency distribution of local correlation values for the same comparisons shown in (A).

**Appendix Figure S4. Features of nucleosome positioning around active promoters in clone 1-2**. (**A**) Plot of interphase and metaphase nucleosome occupancy scores, averaged at the TSSs of 15,073 expressed protein genes. (**B**) Linker length distributions near TSSs showing that shorter linkers are further enriched in promoter regions during interphase. (**C**) Histogram showing the difference in unique nucleosome counts within ±1000 bp of each TSS between interphase and metaphase, for genes in the top and bottom 50% of expression levels.

**Appendix Figure S5. Nucleosome organization at active enhancers during mitosis in clone 1-2.** (**A**) Linker length distributions near active enhancers in interphase versus metaphase. (**B**) Histogram showing the difference in unique nucleosome counts within ±1000 bp of each active enhancer between interphase and metaphase. On average, interphase exhibits 0.455 more unique nucleosomes per enhancer than metaphase.

**Appendix Figure S6. Nucleosome positioning at CTCF binding sites during mitosis in clone 1-2**. (**A**) Comparison of linker length distributions near CTCF binding sites in interphase and metaphase genomes.

(**B**) Nucleosome occupancy scores at CTCF binding sites from interphase and metaphase chemical maps, with nucleosome occupancy scores further divided into quartiles based on CTCF ChIP-seq signals.

**Appendix Figure S7. Nucleosome positioning at exon-intron and intron-exon junctions in clone 1-2.** (**A**) Nucleosome occupancy scores from chemical maps of interphase (upper panels) and metaphase (lower panels), averaged at exon centers, exon-intron junctions, and intron-exon junctions. A total of 164,916 internal exons from 19,166 protein-coding genes were analyzed and grouped into quartiles based on gene expression levels (FPKM). 22,916 exons from non-expressed genes were excluded. (**B**) Same as panel (A) but using MNase-derived nucleosome maps. (**C**) NCP scores from interphase (red) and metaphase (blue) chemical maps at exon-intron and intron-exon junctions (upper panels), alongside NET-seq data averaged at the same positions (lower panels). (**D**) Chemical NCP scores from interphase (upper panels) and metaphase (lower panels) averaged at exon centers and junctions, stratified into expression-level quartiles based on gene FPKM values.

**Appendix Figure S8. Chemical mapping in independent H4S47C-expressing HeLa S3 cells (clone 1-2) verifies the relationship between DNA cyclizability and nucleosome repositioning during mitosis.** (**A**) Average predicted DNA cyclizability scores (C-scores) for unique nucleosomes and their flanking regions derived from genome-wide interphase and metaphase chemical maps in clone 1-2. (**B**) Zoomed-in view of average genome-wide C-scores within the nucleosome region, comparing interphase and metaphase, corresponding to (A). (**C, D**) Interphase and metaphase nucleosomal C-scores within ±500 bp of transcription start sites (C) and active enhancers (D). (**E**) Nucleosomal C-scores within ±500 bp of CTCF binding sites in interphase and metaphase. (**F**) Same as (E), but for unique nucleosomes located at least 60 bp from the CTCF motif center. (**G**) Average nucleosomal C-scores across chromosomal regions ranked by Giemsa staining patterns.

**Appendix Figure S9. DNA cyclizability scores across individual human chromosomes.** Comparison of nucleosomal C-scores between interphase and metaphase across all human chromosomes.

##### Appendix Table S1: Correlation summaries within each Chromatin State

| | | Corr⩽0.15 | Corr>0.15 | odds | odds ratio | Percentage | Function | p-value |
|---|---|---|---|---|---|---|---|---|
| E1 | ROI | 3783 | 12238 | 30.91% | 6.546 | 0.58% | Active TSS | 0 |
| | BG | 123353 | 2612263 | 4.72% | | | | |
| E2 | ROI | 1018 | 4802 | 21.20% | 4.404 | 0.21% | Flanking TSS | 1E-289 |
| | BG | 126118 | 2619699 | 4.81% | | | | |
| E3 | ROI | 4794 | 12676 | 37.82% | 8.074 | 0.64% | Flanking TSS Upstream | 0 |
| | BG | 122342 | 2611825 | 4.68% | | | | |
| E4 | ROI | 1944 | 7262 | 26.77% | 5.596 | 0.33% | Flanking TSS Downstream | 0 |
| | BG | 125192 | 2617239 | 4.78% | | | | |
| E5 | ROI | 2723 | 69266 | 3.93% | 0.807 | 2.62% | Strong transcription | 5.69E-29 |
| | BG | 124413 | 2555235 | 4.87% | | | | |
| E6 | ROI | 14530 | 314524 | 4.62% | 0.948 | 11.96% | Weak transcription | 2.07E-09 |
| | BG | 112606 | 2309977 | 4.87% | | | | |
| E7 | ROI | 472 | 6560 | 7.20% | 1.487 | 0.26% | Genic enhancer1 | 3.44E-15 |
| | BG | 126664 | 2617941 | 4.84% | | | | |
| E8 | ROI | 265 | 2136 | 12.41% | 2.564 | 0.09% | Genic enhancer2 | 1.34E-37 |
| | BG | 126871 | 2622365 | 4.84% | | | | |
| E9 | ROI | 4359 | 17429 | 25.01% | 5.311 | 0.79% | Active Enhancer 1 | 0 |
| | BG | 122777 | 2607072 | 4.71% | | | | |
| E10 | ROI | 628 | 4807 | 13.06% | 2.705 | 0.20% | Active Enhancer 2 | 6.01E-95 |
| | BG | 126508 | 2619694 | 4.83% | | | | |
| E11 | ROI | 7461 | 52741 | 14.15% | 3.040 | 2.19% | Weak Enhancer | 0 |
| | BG | 119675 | 2571760 | 4.65% | | | | |
| E12 | ROI | 156 | 3073 | 5.08% | 1.048 | 0.12% | ZNF genes & repeats | 0.557024 |
| | BG | 126980 | 2621428 | 4.84% | | | | |
| E13 | ROI | 1020 | 38312 | 2.66% | 0.546 | 1.43% | Heterochromatin | 3.14E-97 |
| | BG | 126116 | 2586189 | 4.88% | | | | |
| E14 | ROI | 29 | 293 | 9.90% | 2.043 | 0.01% | Bivalent/Poised TSS | 0.000735 |
| | BG | 127107 | 2624208 | 4.84% | | | | |
| E15 | ROI | 18 | 265 | 6.79% | 1.402 | 0.01% | Bivalent Enhancer | 0.15662 |
| | BG | 127118 | 2624236 | 4.84% | | | | |
| E16 | ROI | 1317 | 38009 | 3.46% | 0.712 | 1.43% | Repressed PolyComb | 8.42E-37 |
| | BG | 125819 | 2586492 | 4.86% | | | | |
| E17 | ROI | 13046 | 335263 | 3.89% | 0.781 | 12.66% | Weak Repressed PolyComb | 3.6E-161 |
| | BG | 114090 | 2289238 | 4.98% | | | | |
| E18 | ROI | 69573 | 1704845 | 4.08% | 0.652 | 64.49% | Quiescent/Low | 0 |
| | BG | 57563 | 919656 | 6.26% | | | | |

##### Appendix Table S2: Linker Length ($l$) Comparisons Among Chromatin States

| | Interphase | | Metaphase | | odd ratio | Test p-value | | |
|---|---|---|---|---|---|---|---|---|
| | $\#(l \leqslant 30)$ | $\#(l > 30)$ | $\#(l \leqslant 30)$ | $\#(l > 30)$ | | vs E5-E6 | vs E13 | vs E18 |
| E1-E4 | 57230 | 87033 | 40225 | 104294 | 1.705 | 0 | 0 | 0 |
| E5-E6 | 582841 | 960381 | 550112 | 1075258 | 1.186 | -- | -- | -- |
| E9-E11 | 108386 | 166771 | 82769 | 200619 | 1.575 | 0 | 0 | 0 |
| E13 | 56260 | 96007 | 55364 | 101836 | 1.078 | -- | -- | -- |
| E18 | 2341812 | 4613845 | 2147515 | 5134842 | 1.214 | -- | -- | -- |

All comparisons yielded highly significant results (p-values ≈ 0), indicating that interphase enrichment was significantly stronger in E1–E4 and E9–E11 compared to E5–E6, E13, or E18. The odds ratios were 1.705 for E1–E4 and 1.575 for E9–E11, substantially higher than those observed for E5–E6 (1.186), E13 (1.078), and E18 (1.214). These findings demonstrate that E1–E4 and E9–E11 exhibit markedly greater enrichment gains during the transition from metaphase to interphase.

##### Appendix Table S3: Linker Length ($l$) Comparisons for TSS, Active Enhancer, and CTCF

| | Interphase | | Metaphase | | odd ratio | Test vs rest Whole Genome region | |
|---|---|---|---|---|---|---|---|
| | $\#(l \leqslant 30)$ | $\#(l > 30)$ | $\#(l \leqslant 30)$ | $\#(l > 30)$ | | odd ratio (other region) | p-value |
| TSS | 48453 | 79791 | 36025 | 93094 | 1.569 | 1.225 | 0 |
| Active Enhancer | 26854 | 39759 | 19893 | 49023 | 1.664 | 1.226 | 0 |
| CTCF | 219217 | 305247 | 153377 | 379739 | 1.778 | 1.205 | 0 |
| Whole | 3716677 | 7232156 | 3374368 | 8063849 | 1.228 | -- | -- |

All comparisons yielded highly significant results (p-values ≈ 0), indicating that short linker length enrichment in interphase was significantly stronger in TSS, active enhancer, and CTCF regions compared to genome-wide background regions with those elements excluded. Specifically, the odds ratios for interphase over metaphase were 1.569 for TSSs, 1.664 for active enhancers, and 1.778 for CTCF sites, each notably higher than the corresponding background odds ratios of 1.225, 1.226, and 1.205 (1.228 for the whole genome). These findings demonstrate that these regulatory regions exhibit markedly greater gains in short linker length during the transition from metaphase to interphase compared to the rest of the genome.

### Figure S1

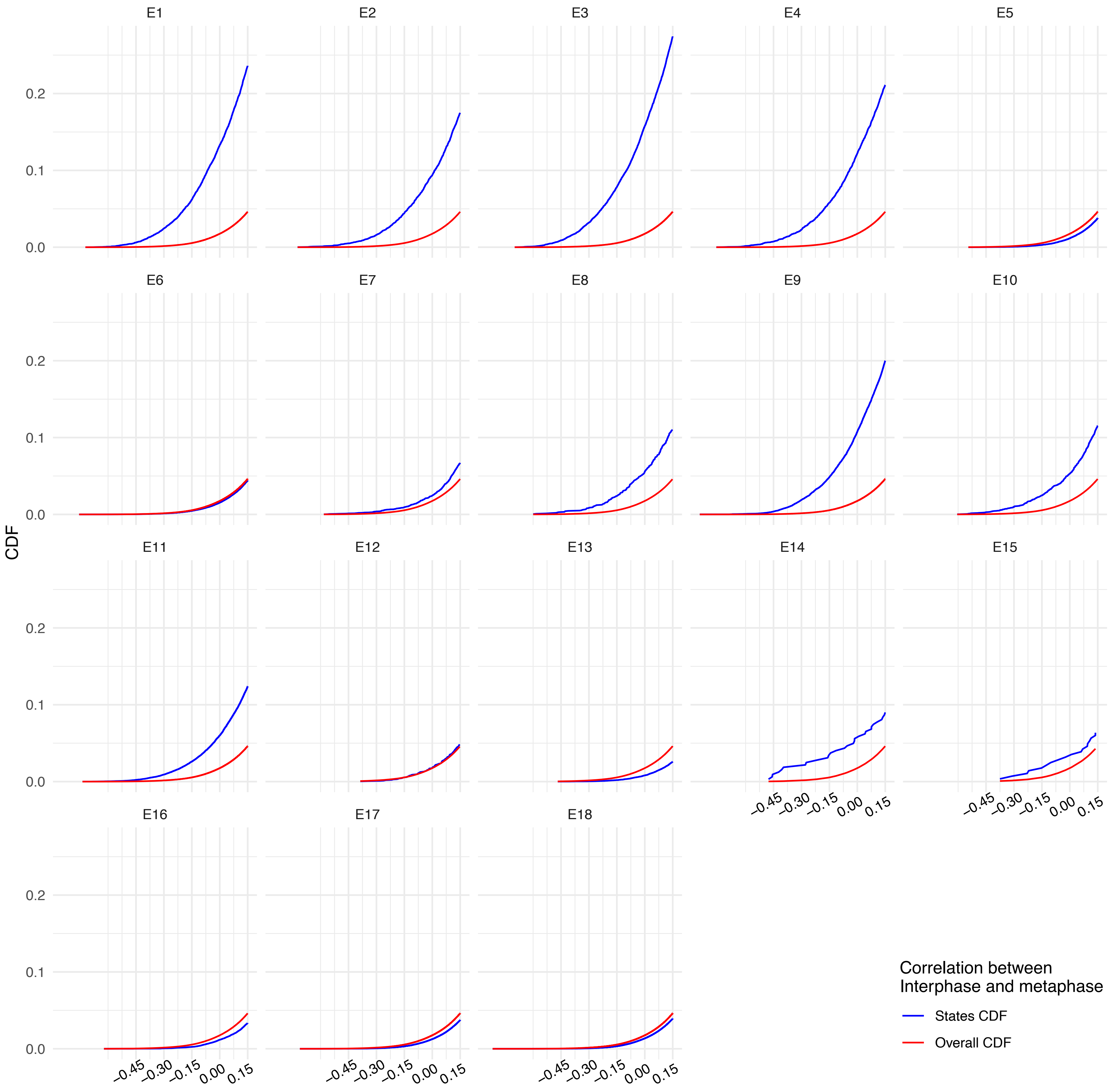
E1
E2
E3
E4
E5
E6
E7
E8
E9
E10
E11
E12
E13
E14
E15
E16
E17
E18
CDF
0.0
0.1
0.2
−0.45
−0.30
−0.15
0.00
0.15
Correlation between
Interphase and metaphase
States CDF
Overall CDF

### Figure S2

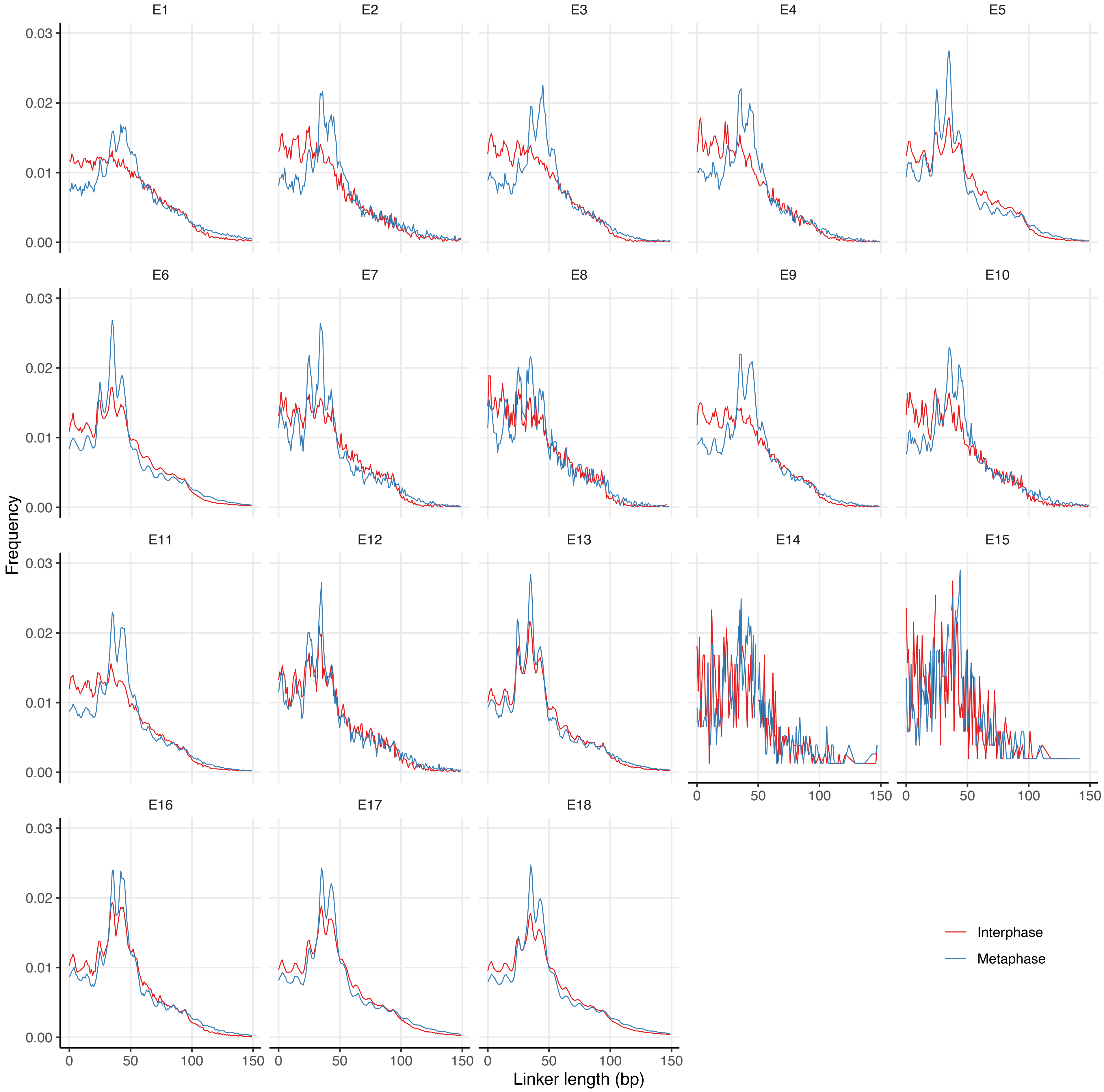
E1
E2
E3
E4
E5
E6
E7
E8
E9
E10
E11
E12
E13
E14
E15
E16
E17
E18
0.00
0.01
0.02
0.03
0
50
100
150
Frequency
Linker length (bp)
Interphase
Metaphase

### Figure S3

A

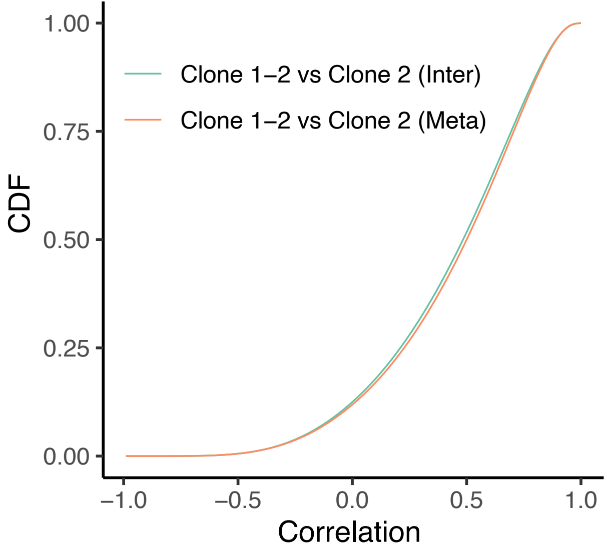

CDF
1.00
0.75
0.50
0.25
0.00
Clone 1−2 vs Clone 2 (Inter)
Clone 1−2 vs Clone 2 (Meta)
−1.0
−0.5
0.0
0.5
1.0
Correlation


B

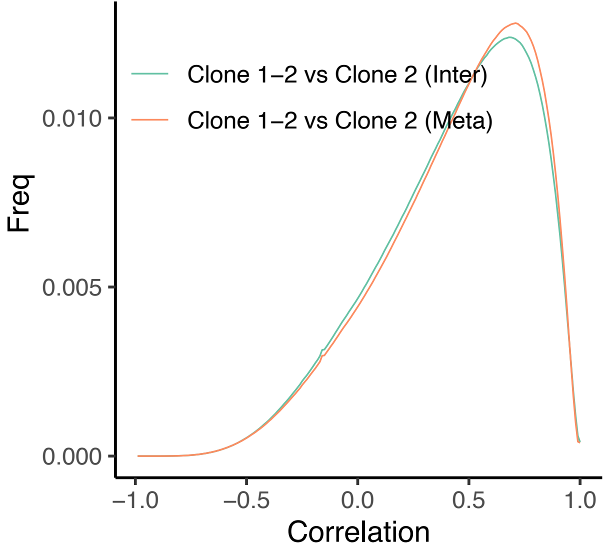

Freq
0.010
0.005
0.000
Clone 1−2 vs Clone 2 (Inter)
Clone 1−2 vs Clone 2 (Meta)
−1.0
−0.5
0.0
0.5
1.0
Correlation

### Figure S4

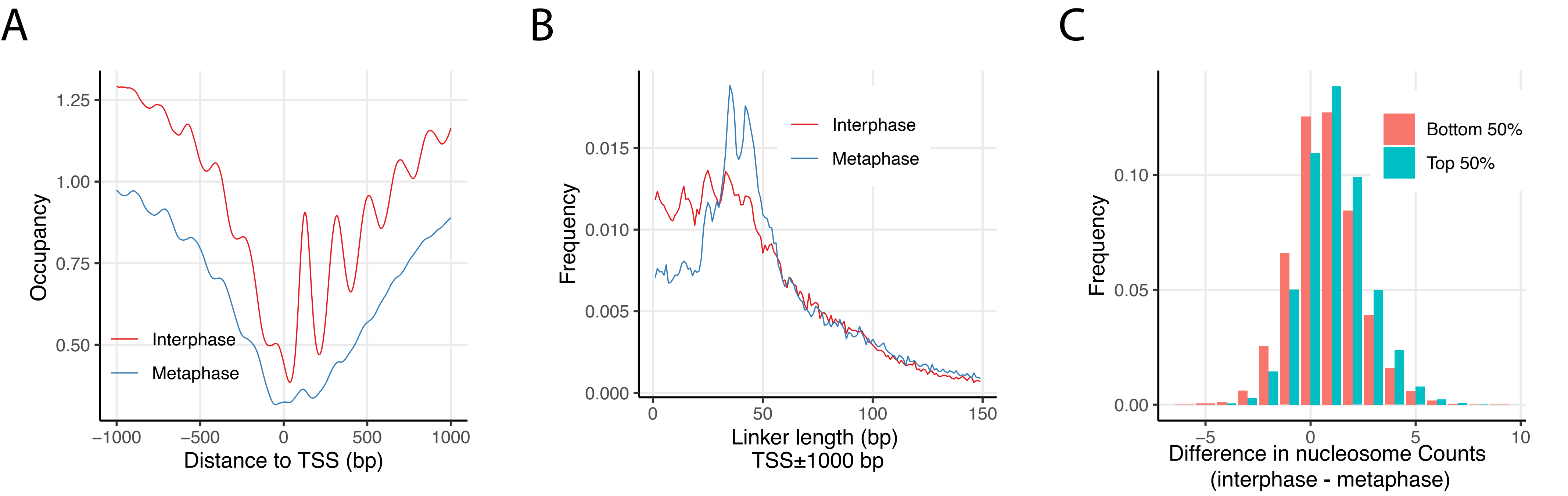
A
B
C
Occupancy
1.25
1.00
0.75
0.50
Interphase
Metaphase
−1000
−500
0
500
1000
Distance to TSS (bp)
Frequency
0.015
0.010
0.005
0.000
Interphase
Metaphase
0
50
100
150
Linker length (bp)
TSS±1000 bp
Frequency
0.10
0.05
0.00
Bottom 50%
Top 50%
−5
0
5
10
Difference in nucleosome Counts
(interphase - metaphase)

### Figure S5

A

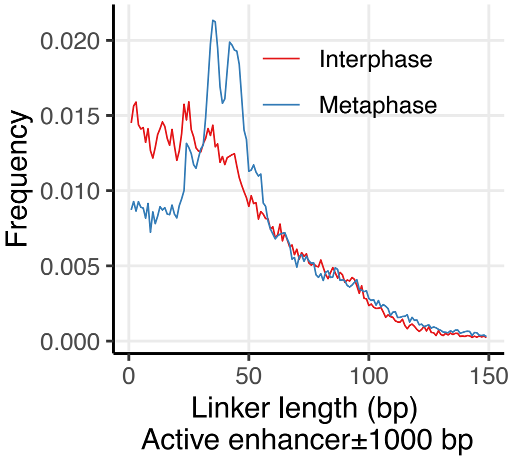
0.020
0.015
0.010
0.005
0.000
Frequency
Interphase
Metaphase
0
50
100
150
Linker length (bp)
Active enhancer±1000 bp

B

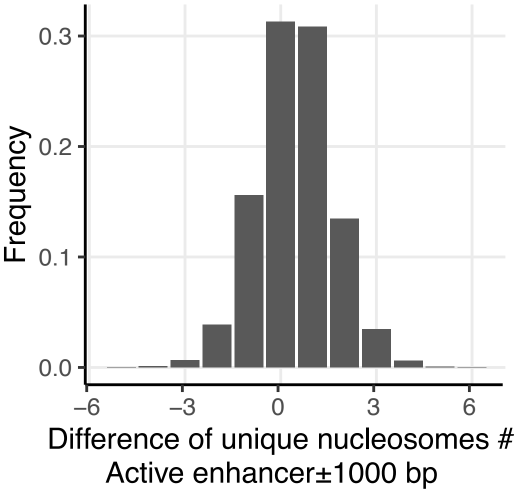
0.3
0.2
0.1
0.0
Frequency
–6
–3
0
3
6
Difference of unique nucleosomes #
Active enhancer±1000 bp

### Figure S6

A

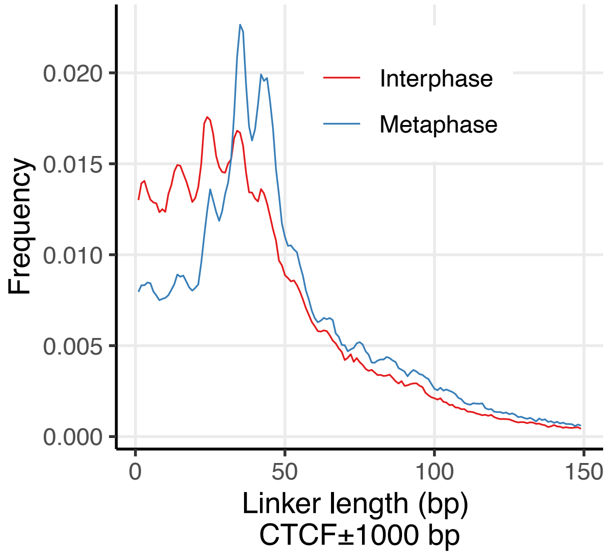

0.020
0.015
0.010
0.005
0.000
Frequency
Interphase
Metaphase
0
50
100
150
Linker length (bp)
CTCF±1000 bp


B

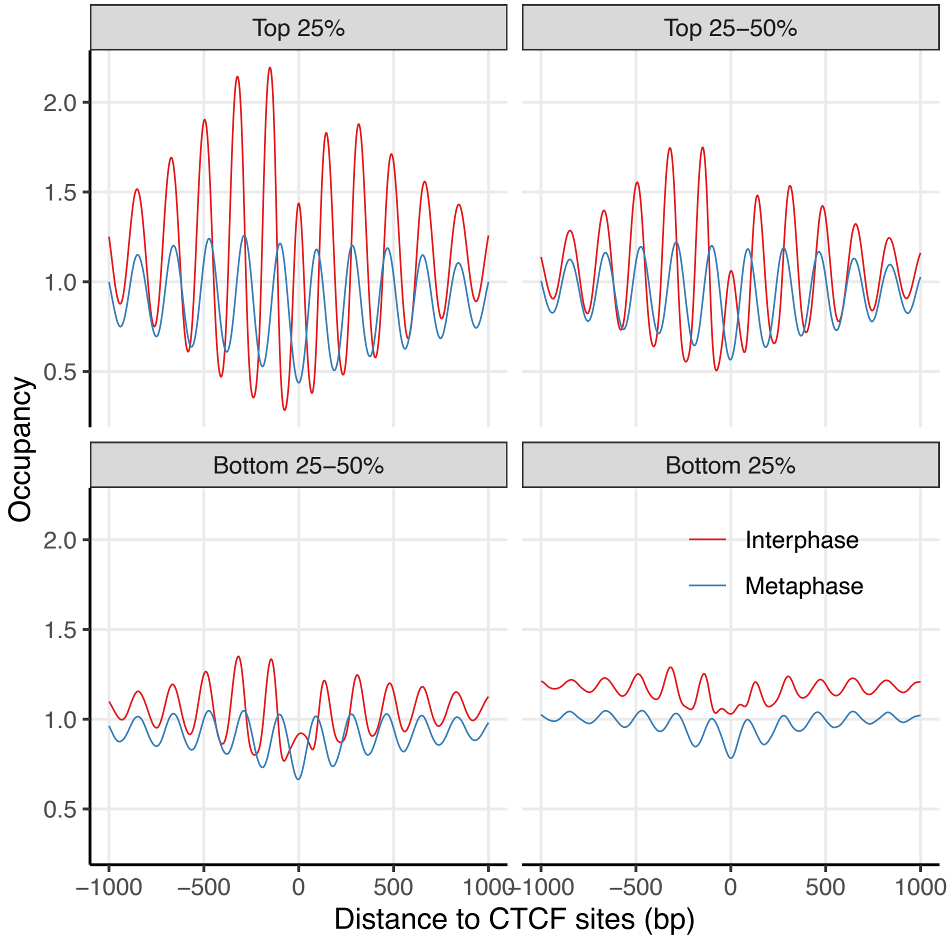

Top 25%
Top 25–50%
Bottom 25–50%
Bottom 25%
2.0
1.5
1.0
0.5
Occupancy
Interphase
Metaphase
−1000
−500
0
500
1000
Distance to CTCF sites (bp)

### Figure S7

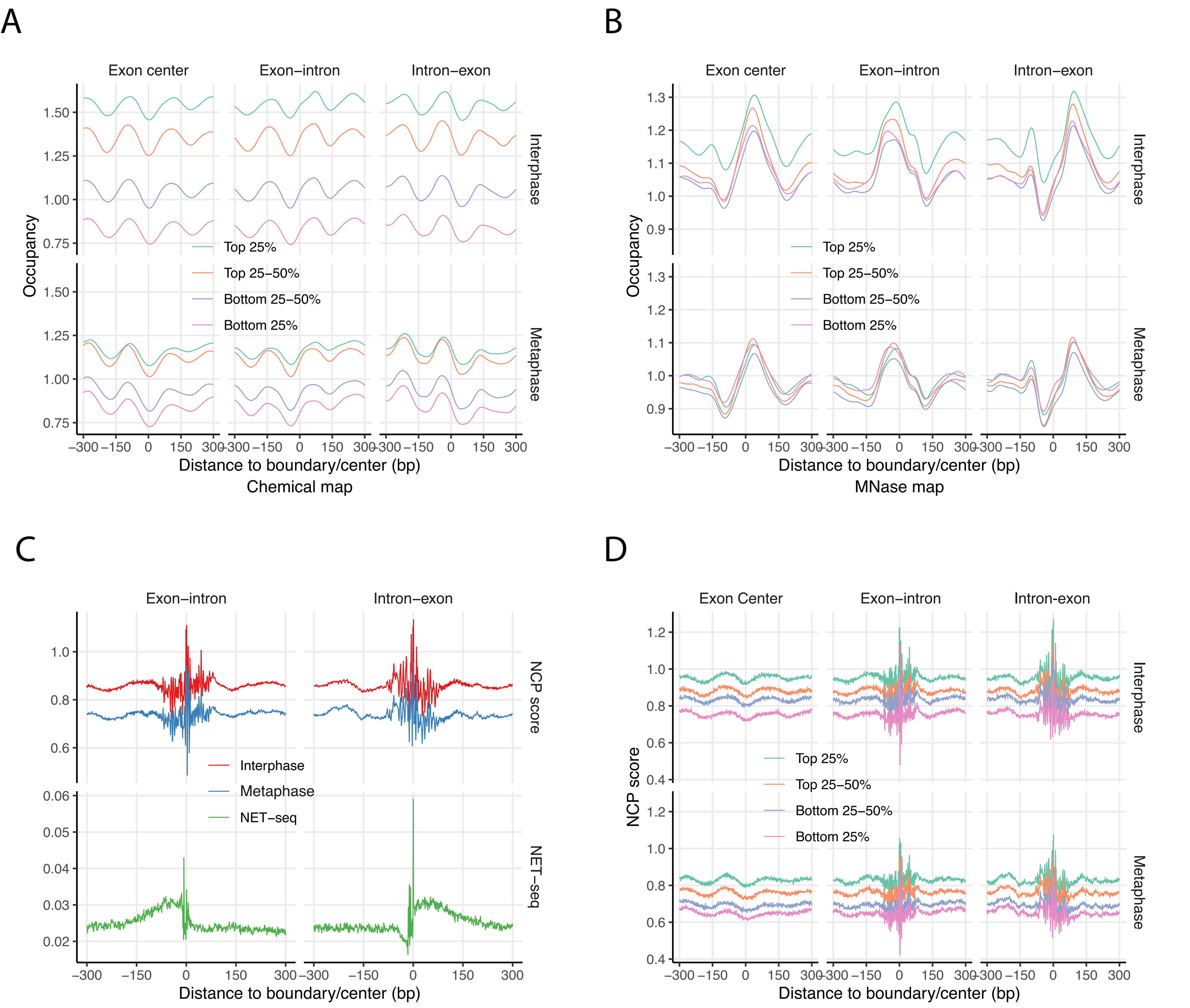
A
Exon center
Exon−intron
Intron−exon
Interphase
Metaphase
Occupancy
1.50
1.25
1.00
0.75
Top 25%
Top 25−50%
Bottom 25−50%
Bottom 25%
−300 −150 0 150 300
Distance to boundary/center (bp)
Chemical map
B
Exon center
Exon−intron
Intron−exon
Interphase
Metaphase
Occupancy
1.3
1.2
1.1
1.0
0.9
Top 25%
Top 25−50%
Bottom 25−50%
Bottom 25%
−300 −150 0 150 300
Distance to boundary/center (bp)
MNase map
C
Exon−intron
Intron−exon
NCP score
NET−seq
1.0
0.8
0.6
0.06
0.05
0.04
0.03
0.02
Interphase
Metaphase
NET−seq
−300 −150 0 150 300
Distance to boundary/center (bp)
D
Exon Center
Exon−intron
Intron−exon
Interphase
Metaphase
NCP score
1.2
1.0
0.8
0.6
0.4
Top 25%
Top 25−50%
Bottom 25−50%
Bottom 25%
−300 −150 0 150 300
Distance to boundary/center (bp)

### Figure S8

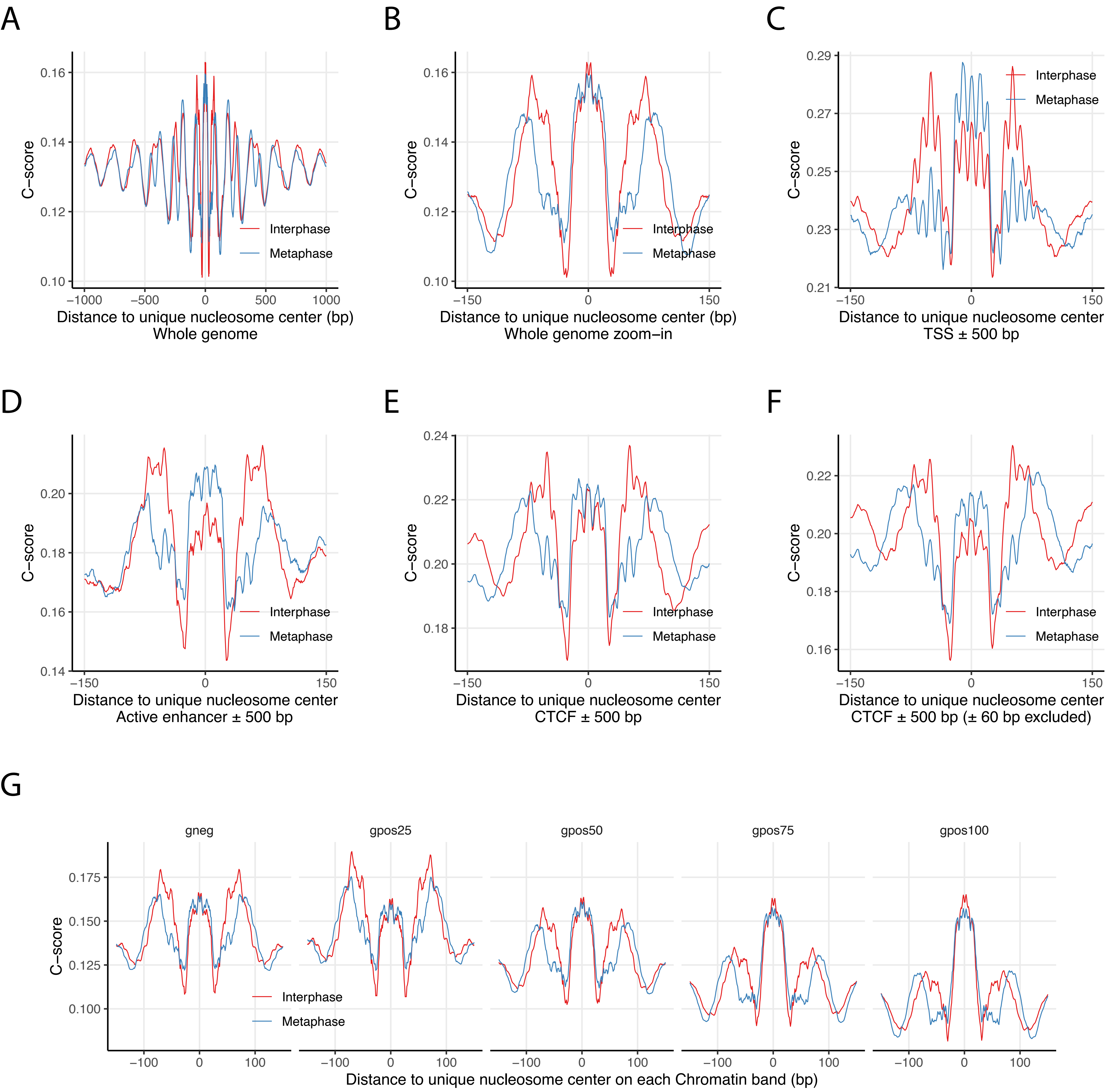
A
B
C
D
E
F
G
C−score
Interphase
Metaphase
Distance to unique nucleosome center (bp)
Whole genome
Whole genome zoom−in
Distance to unique nucleosome center
TSS ± 500 bp
Active enhancer ± 500 bp
CTCF ± 500 bp
CTCF ± 500 bp (± 60 bp excluded)
gneg
gpos25
gpos50
gpos75
gpos100
Distance to unique nucleosome center on each Chromatin band (bp)

### Figure S9

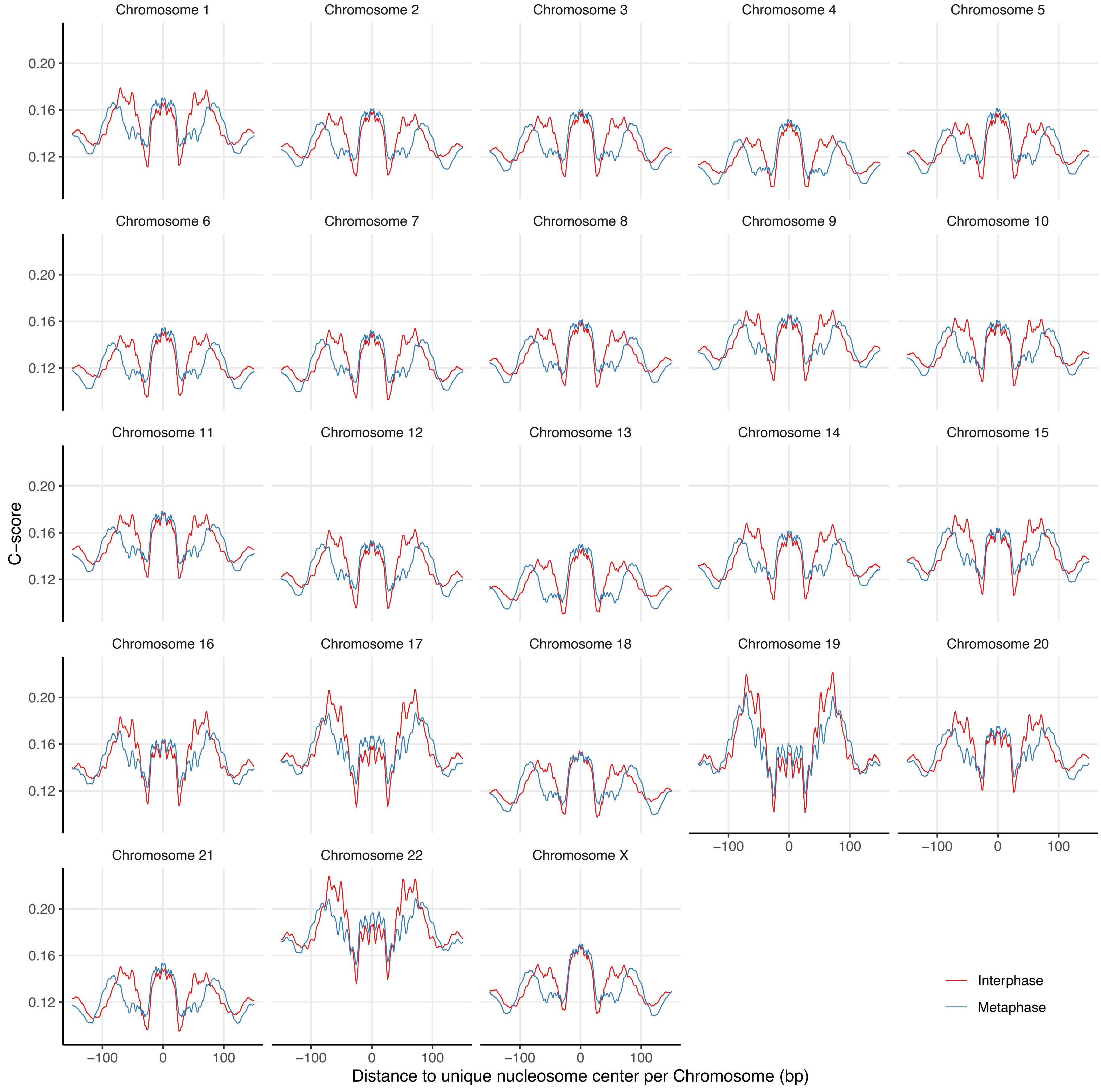

Chromosome 1
Chromosome 2
Chromosome 3
Chromosome 4
Chromosome 5
Chromosome 6
Chromosome 7
Chromosome 8
Chromosome 9
Chromosome 10
Chromosome 11
Chromosome 12
Chromosome 13
Chromosome 14
Chromosome 15
Chromosome 16
Chromosome 17
Chromosome 18
Chromosome 19
Chromosome 20
Chromosome 21
Chromosome 22
Chromosome X
0.20
0.16
0.12
C−score
−100
0
100
Distance to unique nucleosome center per Chromosome (bp)
Interphase
Metaphase
